# Circulating MicroRNAs and Their Associations with Neurotrophic, Inflammatory and Glutamate Markers in Healthy Volunteers

**DOI:** 10.64898/2026.08.04.742737

**Authors:** Aedan O’Shea, Natasha L Mason, Jacco J Briede, Rudy Schreiber, Marcha C.T. Verheijen, Julian Krauskopf, Johannes G Ramaekers

## Abstract

Psilocybin acutely alters neurotrophic, neurochemical, and immune markers, but the relationships between these responses and circulating microRNAs (miRNAs), i.e. non-coding RNAs that regulate post-transcriptional gene expression, remain unclear. In a randomized, double-blind, placebo-controlled study of 62 healthy adults who received psilocybin (0.17 mg/kg) or placebo, we previously demonstrated that let-7g-5p and miR-150-5p were transiently differentially expressed 360 minutes after psilocybin administration. Here, we examined whether changes in these miRNAs were associated with concurrent neurotrophic, inflammatory, pharmacokinetic, and glutamatergic measures. Expression changes from baseline to 360 min and 7 days were analysed using linear regression against changes in BDNF, TNF-α, IL-6, C-reactive protein, cortisol, medial prefrontal cortex glutamate/total creatine, and psilocin concentrations. Psilocybin increased let-7g-5p and decreased miR-150-5p expression. Changes in let-7g-5p were positively associated with psilocin concentrations, suggesting sensitivity to inter-individual pharmacokinetic variability, whereas miR-150-5p showed no concentration-dependent association. In both groups, miRNA changes were negatively related to baseline expression: lower baseline let-7g-5p predicted larger increases, whereas higher baseline miR-150-5p predicted larger decreases. BDNF changes were associated with both miRNAs under placebo but not psilocybin, consistent with reduced between-subject variability and a flattened BDNF-miRNA relationship following treatment. Medial prefrontal glutamate was negatively associated with miR-150-5p change under psilocybin. No associations were found with immune biomarkers. Together, these findings support the predicted involvement of let-7g-5p and miR-150-5p in neuroplasticity and their potential as accessible blood-based biomarkers of individual neurobiological responsiveness to psilocybin and other psychedelics.

## Introduction

Interest in the therapeutic potential of psychedelic drugs for psychiatric disorders, such as Major Depressive Disorder (MDD), has sharply increased in recent years, with psilocybin gaining particular attention (Nutt et al., 2020). Psilocybin (O-phosphoryl-4-hydroxy-N,N-dimethyltryptamine) is a psychedelic drug naturally occurring in psychedelic mushrooms (Meyer & Slot, 2023). Psilocybin and its dephosphorylated pharmacologically active metabolite, psilocin (4-hydroxy-N,N-dimethyltryptamine), were identified as the compounds responsible for the subjective psychedelic effects of these mushrooms, including changes in mood, cognition, and sensory perception (Hofmann et al., 1958; Jaster et al., 2022; Mason et al., 2020; Nutt et al., 2023). Structurally, both compounds share a tryptamine scaffold, making psilocybin a serotonergic classical psychedelic (Jaster et al., 2022). Psilocin acts as a non-selective partial agonist of the serotonin-2A receptor (5HT-2AR) (Mason et al., 2020; Olson, 2022).

At the systems level, psilocybin enhances global neural connectivity, and produces region dependent alterations in glutamate, increasing concentrations in the medial prefrontal cortex (mPFC) and decreasing them in the hippocampus, which drive downstream neuroplasticity (Girn et al., 2026; Mason et al., 2020; Vollenweider & Kometer, 2010). At the molecular level, activation of downstream targets including tropomyosin receptor kinase B (TrkB) which is the primary target for brain-derived neurotrophic factor (BDNF), along with mTOR signalling and AMPA-type glutamate receptor engagement, may be necessary to produce sustained neuroplastic effects (Jaster et al., 2022; Olson, 2022). Preclinical evidence supports elevated BDNF levels and increased dendritic spine density following psilocybin (Ly et al., 2018). However, peripheral BDNF measurements in humans yield inconsistent findings (Calder et al., 2025). This inconsistency points to a broader need for molecular intermediaries that can capture upstream regulatory activity before it manifests at the protein level.

A potential intermediary for studying biological effects of psilocybin or other psychedelics upstream of measurable protein changes are micro RNAs (miRNAs), which are short, non-coding RNAs that regulate gene expression at the post-transcription level, thereby influencing a broad range of cellular processes (Bartel, 2004). When present in blood and other biological fluids, these circulating miRNAs are relatively stable and can be measured non-invasively, making them promising peripheral markers of biological responses to psilocybin (Mitchell et al., 2008; Sohel, 2016).

We have previously demonstrated that psilocybin produces transient alterations in circulating miRNA expression in a sample of healthy volunteers, supporting an acute gene-regulatory response without sustained miRNA changes at 7 days (O’Shea et al., 2026). The miRNAs which were altered by psilocybin, let-7g-5p and miR-150-5p, were found to potentially influence molecular processes involved in neuroplasticity (e.g., TrkA, MAPK), inflammation (e.g., Interleukins, TGF-β), and transcriptional regulation (e.g., RNA polymerase II, SMAD2/3/4) when assessed by over-representation analysis (ORA). As comparatively stable markers of upstream gene regulation, circulating miRNAs may capture subtle molecular changes that are difficult to detect through downstream protein concentrations, such as BDNF, which is limited by measurement-related variability (Calder et al., 2025). Taken together, miRNA expression levels could grant insight to molecular mechanisms of psilocybin, though empirical evidence for direct protein measures reflecting these changes have not been previously studied.

In the current study, we examined whether changes in the expression of these specific miRNAs are associated with parallel alterations in key immune markers (IL-6, TNFα, CRP, cortisol) observed acutely or seven days after psilocybin administration in the same study, as well as with the neurotrophic factor BDNF and medial prefrontal cortex (mPFC) glutamate/total creatine (tCr) measured concurrently. These analyses aim to (1) improve understanding of the neurobiological effects of psilocybin and (2) evaluate the potential of the miRNAs let-7g-5p and miR-150-5p as biomarkers for psilocybin-induced inflammatory and neurotrophic changes.

## Materials and Methods

### Participants and Study design

Plasma circulating miRNA expression profiles were assessed in the blood plasma of 62 healthy adult participants as part of a randomised, placebo-controlled, double-blind, parallel-group design (O’Shea et al., 2026; Mason et al., 2020, 2023). Inclusion and exclusion criteria, ethical approval, and full study procedures are described in detail elsewhere (O’Shea et al., 2026). Participants were allocated to treatment groups and received either a single oral dose of psilocybin (0.17 mg/kg; n=31) or placebo (n=31). For the present analysis, participants with no detectable change in either miRNA (both delta scores = 0) were excluded, as these values likely reflect failed or missing miRNA quantification rather than a true biological response, resulting in a final analytic sample of N= 50 (Psilocybin; n=22, Placebo; n=28).

### Micro-RNA and Marker Delta scores

Blood samples for circulating miRNA analysis were obtained at baseline, 360 minutes post-administration, and 7 days post-administration. Variance stabilizing transformation (VST) miRNA expression values for let-7g-5p and miR-150-5p were obtained from the prior circulating miRNA analysis (O’Shea et al, 2026). Delta scores were calculated as the difference between expression at 360 minutes and baseline for each participant. These delta scores served as the predictor variables in all regression models reported.

Biomarker outcome variables were selected based on biological relevance, temporal proximity to the 6-hour circulating miRNA sampling point, and prior evidence of psilocybin-related modulation in the study sample (Mason et al, 2023): i.e.: a single dose of psilocybin acutely increased glutamate concentration in the medial prefrontal (mPFC) cortex and cortisol levels and decreased TNF-α. Sub-acutely, IL-6 and CRP decreased. In addition, we included BDNF changes in this cohort that have not been previously reported. Change in BDNF, TNF-α, IL-6, and psilocin concentration were included for analysed against baseline-to-360-minute change scores in miRNA. A second IL-6 change score, reflecting the 7-day follow-up timepoint, was additionally included. CRP was log10-transformed and analysed as both an acute change score and a long-term change score. Cortisol was analysed as a baseline-to-80-minute change score, reflecting the acute stress-response window. mPFC glutamate concentration was obtained from MRS acquired approximately 65 min post-administration and analysed as a single post-dose measure, as no baseline MRS acquisition was available (Mason et al., 2020). Psilocin concentration was analysed only in the psilocybin group, as placebo participants had no detectable psilocin. Table 1 provides a summary of all associations that were assessed.

**Table 1.** Summary of neurotrophic, inflammatory, pharmacokinetic, and neurochemical outcomes whose associations with changes in miRNA expression were assessed. Δ indicates that the measure is baseline corrected. Time after treatment (min) is provided in brackets.

| Treatment day (psilocybin or placebo) | Day 7 after treatment |
| --- | --- |
| $\Delta$ miRNA (360 min) | $\Delta$ miRNA |
| $\Delta$ CRP (60-120 min) | $\Delta$ CRP |
| $\Delta$ IL-6 (60-120 min) | $\Delta$ IL-6 |
| $\Delta$ TNF-α (60-120 min) | |
| $\Delta$ BDNF (360 min) | |
| $\Delta$ cortisol (80 min) | |
| Psilocin (360 min) |  |
| Glutamate/tCr (60-120 min) |  |

### Statistical analysis

Associations between miRNA delta scores and biomarker outcomes were assessed using simple linear regression, with one model fitted for each miRNA-outcome pair. The two miRNA predictors, let-7g-5p and miR-150-5p, were analysed separately. The biological markers were the outcomes and were assessed separately. This univariate modelling strategy was chosen because the analysis was exploratory and aimed to identify candidate miRNA-biomarker associations rather than estimate independent multivariable effects.

Models were run in two stages. First, primary analyses were conducted in the full analytic sample, pooled across treatment groups, to test overall miRNA– biomarker associations. Second, exploratory subgroup analyses were conducted separately within the placebo and psilocybin groups to assess whether associations were condition specific. Models were fitted using complete cases for each miRNA-outcome pair, the number of observations per model is reported in the results. Nominal statistical significance was defined as p < 0.05. Benjamini–Hochberg false discovery rate correction and Bonferroni correction were applied across models and within each miRNA predictor separately, and adjusted p-values are reported alongside uncorrected p-values for transparency. Given the exploratory nature of the analysis, uncorrected p-values were used as the basis for interpretation, with corrected values provided for reference. All analyses were conducted in R version 4.4.1 using base R functions, including the stats package. Data handling was performed using dplyr version 1.1.4, and scatter plots with group-specific regression lines were generated using ggplot2 version 4.0.0.

## Results

### Participants and Analytical sample

A total of 50 participants were included in the final analytic sample. The final sample comprised 28 placebo and 22 psilocybin participants. Demographic characteristics of the full study cohort are reported elsewhere (O’Shea et al., 2026).

### miRNA expressions following psilocybin administration

Consistent with prior findings from this cohort (O’Shea et al., 2026), let-7g-5p (t=2.05; p=.04) expression increased from baseline to 360 minutes in the psilocybin group, while miR-150-5p expression decreased (t=2.18; p=.037). Both directions of change were also present in the placebo group, though with greater variance and a mean closer to zero (Figure 1A, 1B). Delta scores for both miRNAs showed a strong negative relationship with baseline expression levels in both groups (Figure 2A, 2B): participants with lower baseline let-7g-5p expression showed larger increases, while participants with higher baseline miR-150-5p expression showed larger decreases respectively. These relationships were present in both treatment conditions but were numerically stronger in the psilocybin group (let-7g-5p: placebo R^2^=0.49, p<0.001; psilocybin R^2^=0.59, p<0.001; miR-150-5p: placebo R^2^=0.49, p<0.001; psilocybin R^2^=0.64, p<0.001), suggesting that psilocybin amplifies baseline-dependent miRNA regulation.

**Figure 1.**
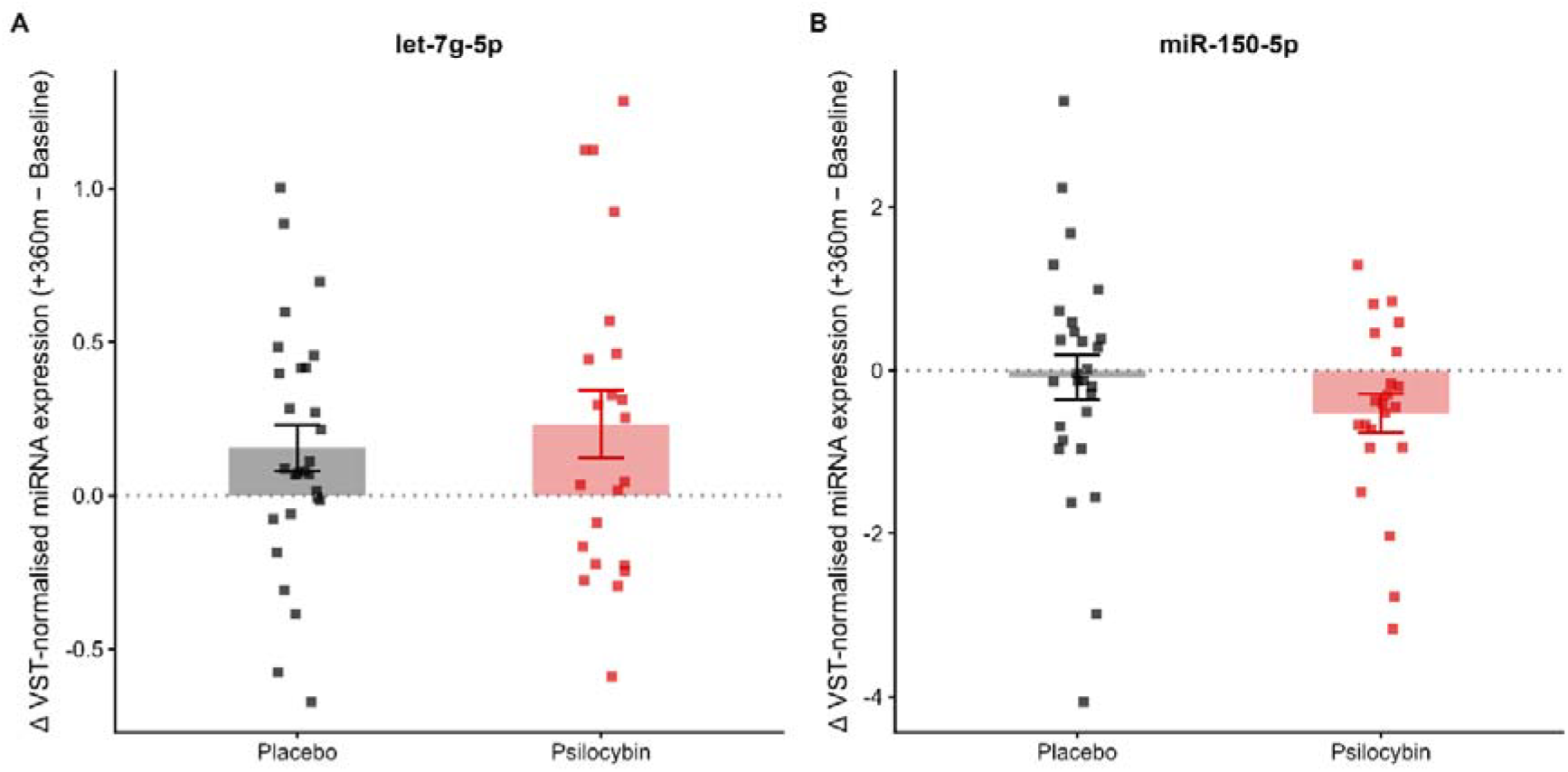
Acute change in circulating miRNA expression following psilocybin or placebo administration. Change in VST-normalised expression of let-7g-5p and miR-150-5p from Baseline to +360 min is shown for placebo and psilocybin participants. Bars indicate group means and error bars indicate SEM; individual participants are shown as square points. Placebo participants are shown in black and psilocybin participants in red. The dotted horizontal line indicates no change from baseline. Values represent +360 min minus Baseline expression.

**Figure 2.**
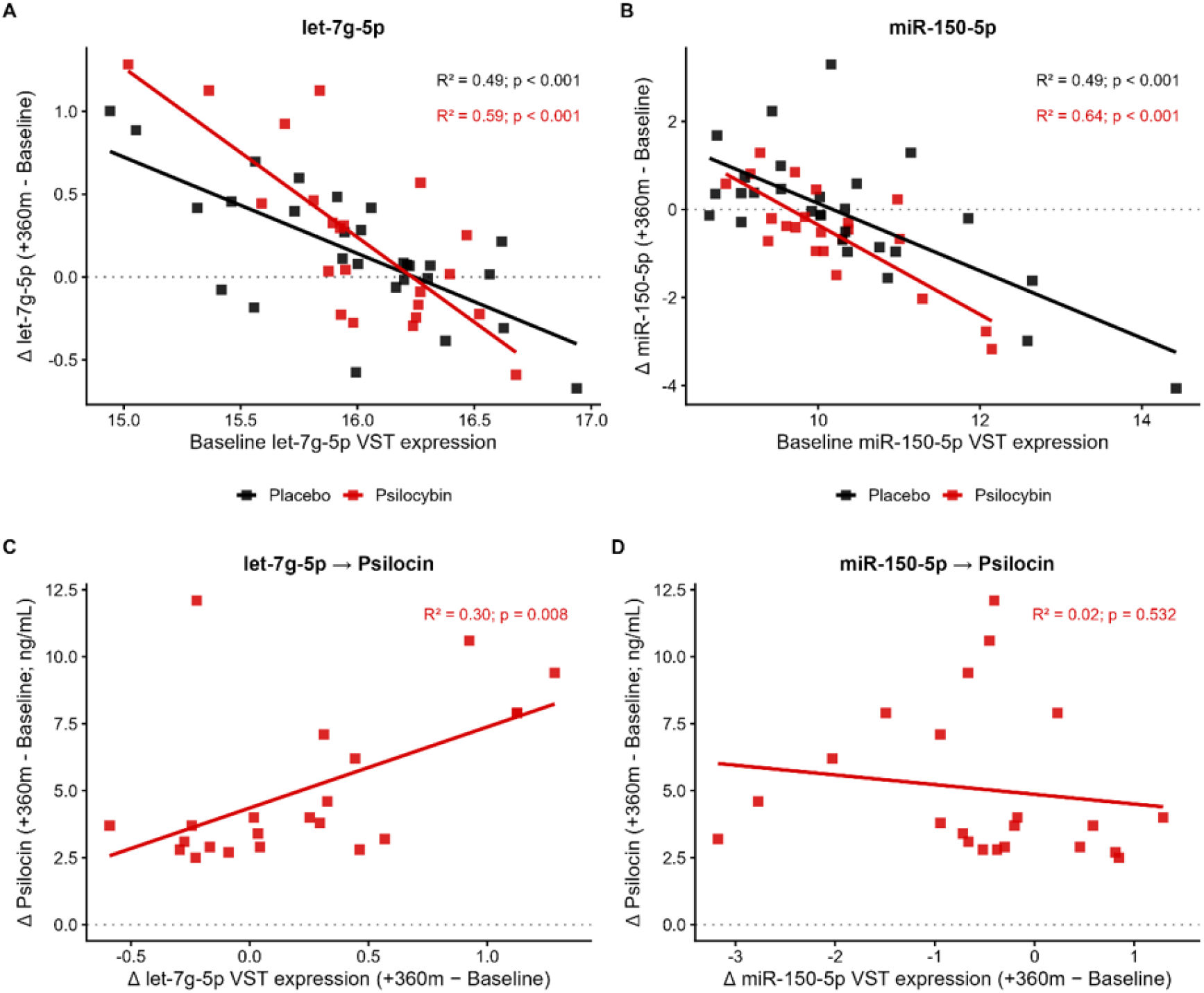
Baseline-dependent miRNA changes and association with psilocin exposure. Panels A and B show baseline VST-normalised expression of let-7g-5p and miR-150-5p plotted against subsequent change in expression from Baseline to +360 min. Placebo participants are shown in black and psilocybin participants in red, with separate ordinary least-squares regression lines by group. Panels C and D show psilocybin-only associations between change in miRNA expression and change in psilocin concentration from Baseline to +360 min. R^2^ and p-values are from the linear regression model shown in each panel and are unadjusted. Baseline-change associations in panels A and B should be interpreted descriptively, because change scores include the baseline value in their calculation.

### Associations between miRNA delta scores and psilocin concentration

Psilocin concentration was assessed only in the psilocybin group, as placebo participants had no detectable psilocin. Within this group, let-7g-5p delta scores were positively associated with psilocin concentration at 360 minutes (R^2^=0.30, p=0.008; Figure 2C), indicating that participants with greater acute increases in let-7g-5p expression also had higher psilocin levels. No significant association was observed between miR-150-5p delta scores and psilocin concentration (R^2^=0.02, p=0.532; Figure 2D).

### miRNA delta scores as predictors of delta BDNF and mPFC glutamate

Mean BDNF change scores were near zero in both treatment groups, with substantial inter-individual variability in both conditions (t<1.8; p=NS) (Figure 3A). mPFC glutamate/tCr values, were higher in the psilocybin group as compared to the placebo group (t=2.52; p=.015) (Figure 3B). In the pooled primary analysis, both miRNA delta scores were nominally significantly associated with BDNF change scores, let-7g-5p (R^2^=0.10, p=0.028) and miR-150-5p (R^2^=0.09, p=0.031).

**Figure 3.**
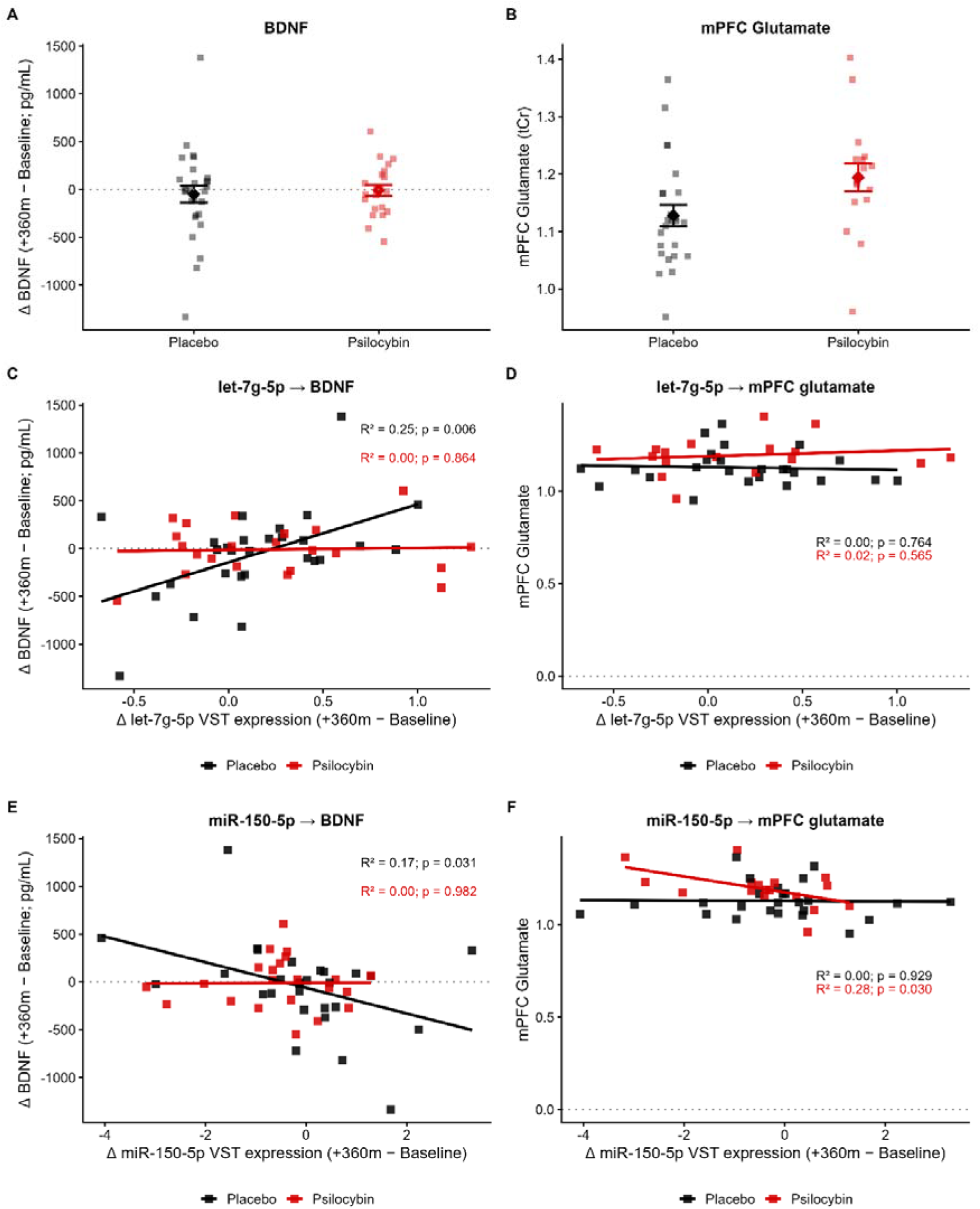
Associations between miRNA expression changes, BDNF change, and mPFC glutamate/tCr. Panels A and B show group-level distributions of BDNF change from Baseline to +360 min and mPFC glutamate/tCr, respectively. Bars indicate group means and error bars indicate SEM; individual participants are shown as square points. Panels C and E show associations between miRNA expression change and BDNF change for let-7g-5p and miR-150-5p. Panels D and F show associations between miRNA expression change and mPFC glutamate/tCr. Placebo participants are shown in black and psilocybin participants in red, with separate regression lines by group. R^2^ and p-values are from group-specific ordinary least-squares regressions and are unadjusted. Analyses involving mPFC glutamate/tCr include fewer participants because of missing MRS data.

Analyses per treatment group revealed that both overall associations were largely attributable to the placebo subgroup (let-7g-5p: placebo R^2^ = 0.25, p=0.006, psilocybin R^2^ = 0.00, p = 0.864; miR-150-5p: placebo R^2^ = 0.17, p = 0.031, psilocybin R^2^ = 0.00, p = 0.982; Figure 3C, 3E). In the psilocybin group, the regression slope was flat for both predictors, thus no evidence of a corresponding miRNA-BDNF association was observed in the psilocybin group.

Values for mPFC glutamate represent glutamate concentration ratios relative to tCr. For mPFC glutamate/tCr, no significant associations were observed in the pooled analysis or in the placebo group for either miRNA predictor (let-7g-5p: placebo R^2^ = 0.00, p = 0.764, psilocybin R^2^ = 0.02, p = 0.565; Figure 3D). However, miR-150-5p delta scores were nominally significantly associated with mPFC glutamate concentration in the psilocybin group specifically (R^2^ = 0.28, p = 0.030), with no corresponding association in placebo (R^2^ = 0.00, p = 0.929; Figure 3F).

### miRNA delta scores as predictors of delta inflammatory and stress markers

No significant associations were found between either miRNA delta score and change scores for TNF-α, IL-6, CRP or cortisol in either primary pooled or group-separated secondary analyses (all p>0.05).

### Correction for multiple comparisons

No miRNA-biomarker associations survived Benjamini-Hochberg correction when applied across all models simultaneously, consistent with the exploratory nature of the analysis, uncorrected p-values are reported as the primary inference as previously specified.

## Discussion

The present analysis reconfirmed increments in let-7g-5p and decrements in miR-150-5p under psilocybin as compared to baseline. Increment in let-7g-5p was positively associated with psilocin concentration, whereas miR-150-5p was not. Changes in miRNAs were negatively associated with baseline expression levels in both groups. Participants with lower baseline let-7g-5p expression showed larger increases, while participants with higher baseline miR-150-5p expression showed larger decreases respectively. Associations between these miRNAs and neurotropic and immune biomarkers practically confirmed the predicted involvement of let-7g-5p and miR-150-5 in in neuroplasticity. Associations between delta BDNF and delta miRNAs were present in the placebo group but absent in the group receiving psilocybin. Glutamate concentrations in the mPFC were negatively associate with change in miR-150-5p in the psilocybin group. Changes in immune biomarkers were not associated with changes in let-7g-5p and miR-150-5.

The clearest biomarker association observed in the present analysis was between acute miRNA change and BDNF change. In the pooled primary models, BDNF change from Baseline to +360 min was nominally associated with delta scores for both let-7g-5p and miR-150-5p. However, group-separated follow-up analyses indicated that these associations were primarily driven by placebo participants , whereas the corresponding relationships were essentially absent in the psilocybin group. This pattern suggests that the apparent pooled miRNA-BDNF associations do not reflect psilocybin specific coupling between miRNA regulation and BDNF response. One possible interpretation is that psilocybin induced baseline dependent miRNA normalization, whereby let-7g-5p increases more in individuals with lower baseline expression and miR-150-5p decreases more in individuals with higher baseline expression, reducing between-subject variability in miRNA state and flattening the association with BDNF change.

The association between miR-150-5p delta scores and mPFC glutamate concentration, observed exclusively in the psilocybin group, is an intriguing addition to this pattern. Psilocybin has previously been shown to produce region-specific alterations in glutamate, including elevated concentrations in the mPFC at approximately 65 minutes post-administration (Mason et al., 2020). Increments in glutamate were highest in individuals that showed a larger decrease in miR-150-5p. This finding raises the possibility that acute changes in miR-150-5p expression covary with prefrontal glutamatergic responses, although due to the exploratory design and temporal structure of the analysis, we cannot infer conclusions about causality or directionality. It should be noted however that miR-150-5p has experimentally supported gene targets in pathways relevant to glutamatergic and synaptic signalling (O’Shea et al., 2026; Cui et al., 2025; Huang et al., 2020). Moreover, psychedelics have been associated with alterations in glutamatergic signalling, which are thought to contribute to sustained neuroplasticity and reductions in depressive symptoms (Vollenweider and Kometer, 2010).

Within the psilocybin group, changes in let-7g-5p expression were associated with psilocin concentrations at 360 minutes, with higher concentrations corresponding to greater increases in expression. This finding suggests that the let-7g-5p response was related to the extent of systemic psilocin exposure and may therefore capture inter-individual pharmacokinetic variability. Rather than representing a uniform miRNA response to a fixed psilocybin dose, let-7g-5p expression may reflect differences in the degree to which participants were biologically engaged by the drug. In contrast to miR-150-5p, let-7g-5p may thus provide an exposure-sensitive molecular readout: participants receiving the same administered dose can achieve markedly different psilocin concentrations, and the concentration-dependent let-7g-5p response may indicate how this variability in systemic exposure is translated into a downstream biological response.

This study provides the first practical demonstration in humans that psychedelic-induced changes in circulating miRNA expression are associated with both a peripheral measure of neurotrophic signalling, BDNF, and a central, brain-derived measure of glutamatergic function. This convergence suggests that circulating miRNAs may provide a minimally invasive molecular readout of biological processes relevant to psychedelic-induced neuroplasticity. Circulating miRNAs may also complement, or potentially provide a more reliable alternative to, peripheral BDNF as a biomarker of psychedelic-induced plasticity. Although BDNF is commonly used as an indirect measure of neuroplasticity, circulating concentrations are highly variable and difficult to interpret (Calder et al., 2025). Moreover, peripheral BDNF concentrations cannot necessarily be assumed to reflect BDNF signalling or plasticity within the brain. By measuring relatively stable regulatory molecules that collectively influence multiple plasticity-related pathways, miRNA profiles may provide a broader and potentially more reproducible molecular readout than a single circulating neurotrophic protein. However, any advantage over BDNF remains to be established. Replication in adequately powered samples, using standardized procedures and temporally matched assessments of miRNA expression, peripheral BDNF, and central indices of neuroplasticity, will be necessary to determine whether these associations are reproducible and biologically meaningful.

This study also comes with limitations that warrant cautionary interpretation of the current findings. First, the group-separated analyses were conducted in modest sample sizes and should be viewed as exploratory follow-ups rather than formal evidence of treatment-by-miRNA interaction. Second, peripheral BDNF is an imprecise proxy for central neuroplasticity-related signalling, with substantial measurement variability reported across human psychedelic studies. The apparent absence of miRNA-BDNF associations in the psilocybin group may therefore reflect true condition-specific decoupling, baseline-dependent miRNA normalization, limited statistical power, or measurement variability. Third, these findings should be interpreted as exploratory and condition-dependent, particularly because the reported p-values are unadjusted and no miRNA-BDNF association survived correction across the full model set.

In conclusion, the predicted involvement of the circulating miRNAs let-7g-5p and miR-150-5p in neuroplasticity and inflammatory regulation, together with their observed associations with BDNF and glutamate, suggests that they may serve as accessible blood-based biomarkers of individual neurobiological responsiveness to psilocybin and other psychedelics.

## References

Bartel, D. P. (2004). MicroRNAs: Genomics, Biogenesis, Mechanism, and Function. Cell, 116(2), 281–297. 10.1016/S0092-8674(04)00045-5

Calder, A. E., Hase, A., & Hasler, G. (2025). Effects of psychoplastogens on blood levels of brain-derived neurotrophic factor (BDNF) in humans: A systematic review and meta-analysis. Molecular Psychiatry, 30(2), 763–776. 10.1038/s41380-024-02830-z

Chang, J., Jiang, T., Shan, X., Zhang, M., Li, Y., Qi, X., Bian, Y., & Zhao, L. (2024). Pro-inflammatory cytokines in stress-induced depression: Novel insights into mechanisms and promising therapeutic strategies. Progress in Neuro-Psychopharmacology and Biological Psychiatry, 131, 110931. 10.1016/j.pnpbp.2023.110931

Cui, S., Yu, S., Huang, H.-Y., Lin, Y.-C.-D., Huang, Y., Zhang, B., Xiao, J., Zuo, H., Wang, J., Li, Z., Li, G., Ma, J., Chen, B., Zhang, H., Fu, J., Wang, L., & Huang, H.-D. (2025). miRTarBase 2025: Updates to the collection of experimentally validated microRNA–target interactions. Nucleic Acids Research, 53(D1), D147–D156. 10.1093/nar/gkae1072

Girn, M., Doss, M. K., Roseman, L., Preller, K. H., Palhano-Fontes, F., Pasquini, L., Barrett, F. S., Mallaroni, P., Mason, N. L., Timmermann, C., McCulloch, D. E., Fisher, P. M., Winston, B. S., Moujaes, F., Muller, F., Liechti, M. E., Vollenweider, F. X., Ramaekers, J. G., Kuypers, K., … Bzdok, D. (2026). An international mega-analysis of psychedelic drug effects on brain circuit function. Nature Medicine, 32(4), 1543–1554. 10.1038/s41591-026-04287-9

Hofmann, A., Heim, R., Brack, A., & Kobel, H. (1958). Psilocybin, ein psychotroper Wirkstoff aus dem mexikanischen RauschpilzPsilocybe mexicana Heim. Experientia, 14(3), 107–109. 10.1007/BF02159243

Huang, H.-Y., Lin, Y.-C.-D., Li, J., Huang, K.-Y., Shrestha, S., Hong, H.-C., Tang, Y., Chen, Y.-G., Jin, C.-N., Yu, Y., Xu, J.-T., Li, Y.-M., Cai, X.-X., Zhou, Z.-Y., Chen, X.-H., Pei, Y.-Y., Hu, L., Su, J.-J., Cui, S.-D., … Huang, H.-D. (2020). miRTarBase 2020: Updates to the experimentally validated microRNA-target interaction database. Nucleic Acids Research, 48(D1), D148–D154. 10.1093/nar/gkz896

Jaster, A. M., de la Fuente Revenga, M., & González-Maeso, J. (2022). Molecular targets of psychedelic-induced plasticity. Journal of Neurochemistry, 162(1), 80–88. 10.1111/jnc.15536

Ly, C., Greb, A. C., Cameron, L. P., Wong, J. M., Barragan, E. V., Wilson, P. C., Burbach, K. F., Soltanzadeh Zarandi, S., Sood, A., Paddy, M. R., Duim, W. C., Dennis, M. Y., McAllister, A. K., Ori-McKenney, K. M., Gray, J. A., & Olson, D. E. (2018). Psychedelics Promote Structural and Functional Neural Plasticity. Cell Reports, 23(11), 3170–3182. 10.1016/j.celrep.2018.05.022

Mason, N. L., Kuypers, K. P. C., Müller, F., Reckweg, J., Tse, D. H. Y., Toennes, S. W., Hutten, N. R. P. W., Jansen, J. F. A., Stiers, P., Feilding, A., & Ramaekers, J. G. (2020). Me, myself, bye: Regional alterations in glutamate and the experience of ego dissolution with psilocybin. Neuropsychopharmacology, 45(12), 2003–2011. 10.1038/s41386-020-0718-8

Mason, N. L., Szabo, A., Kuypers, K. P. C., Mallaroni, P. A., de la Torre Fornell, R., Reckweg, J. T., Tse, D. H. Y., Hutten, N. R. P. W., Feilding, A., & Ramaekers, J. G. (2023). Psilocybin induces acute and persisting alterations in immune status in healthy volunteers: An experimental, placebo-controlled study. Brain, Behavior, and Immunity, 114, 299–310. 10.1016/j.bbi.2023.09.004

Meyer, M., & Slot, J. (2023). The evolution and ecology of psilocybin in nature. Fungal Genetics and Biology, 167, 103812. 10.1016/j.fgb.2023.103812

Mitchell, P. S., Parkin, R. K., Kroh, E. M., Fritz, B. R., Wyman, S. K., Pogosova-Agadjanyan, E. L., Peterson, A., Noteboom, J., O’Briant, K. C., Allen, A., Lin, D. W., Urban, N., Drescher, C. W., Knudsen, B. S., Stirewalt, D. L., Gentleman, R., Vessella, R. L., Nelson, P. S., Martin, D. B., & Tewari, M. (2008). Circulating microRNAs as stable blood-based markers for cancer detection. Proceedings of the National Academy of Sciences, 105(30), 10513–10518. 10.1073/pnas.0804549105

Nutt, D., Erritzoe, D., & Carhart-Harris, R. (2020). Psychedelic Psychiatry’s Brave New World. Cell, 181(1), 24–28. 10.1016/j.cell.2020.03.020

Nutt, D., Spriggs, M., & Erritzoe, D. (2023). Psychedelics therapeutics: What we know, what we think, and what we need to research. Neuropharmacology, 223, 109257. 10.1016/j.neuropharm.2022.109257

Olson, D. E. (2022). Biochemical Mechanisms Underlying Psychedelic-Induced Neuroplasticity. Biochemistry, 61(3), 127–136. 10.1021/acs.biochem.1c00812

Sohel, M. H. (2016). Extracellular/Circulating MicroRNAs: Release Mechanisms, Functions and Challenges. Achievements in the Life Sciences, 10(2), 175–186. 10.1016/j.als.2016.11.007

Vollenweider, F. X., & Kometer, M. (2010). The neurobiology of psychedelic drugs: Implications for the treatment of mood disorders. Nature Reviews Neuroscience, 11(9), 642–651. 10.1038/nrn2884

Zeng, Y., Chourpiliadis, C., Hammar, N., Seitz, C., Valdimarsdóttir, U. A., Fang, F., Song, H., & Wei, D. (2024). Inflammatory Biomarkers and Risk of Psychiatric Disorders. JAMA Psychiatry, 81(11), 1118–1129. 10.1001/jamapsychiatry.2024.2185

